# Prospective mapping of viral mutations that escape antibodies used to treat COVID-19

**DOI:** 10.1101/2020.11.30.405472

**Authors:** Tyler N. Starr, Allison J. Greaney, Amin Addetia, William W. Hannon, Manish C. Choudhary, Adam S. Dingens, Jonathan Z. Li, Jesse D. Bloom

**Author notes:** These authors contributed equally to this work (Tyler N. Starr, Allison J. Greaney).

## Abstract

Antibodies are becoming a frontline therapy for SARS-CoV-2, but the risk of viral evolutionary escape remains unclear. Here we map how all mutations to SARS-CoV-2’s receptor-binding domain (RBD) affect binding by the antibodies in Regeneron’s REGN-COV2 cocktail and Eli Lilly’s LY-CoV016. These complete maps uncover a single amino-acid mutation that fully escapes the REGN-COV2 cocktail, which consists of two antibodies targeting distinct structural epitopes. The maps also identify viral mutations that are selected in a persistently infected patient treated with REGN-COV2, as well as in lab viral escape selections. Finally, the maps reveal that mutations escaping each individual antibody are already present in circulating SARS-CoV-2 strains. Overall, these complete escape maps enable immediate interpretation of the consequences of mutations observed during viral surveillance.

---

Antibodies are being developed as therapeutics to combat SARS-CoV-2 (*1*). Antibodies against some other viruses can be rendered ineffective by viral mutations that are selected during treatment of infected patients (*2*, *3*) or that spread globally to confer resistance on entire viral clades (*4*). Therefore, determining *a priori* which SARS-CoV-2 mutations escape key antibodies is essential for assessing how mutations observed during viral surveillance impact the efficacy of antibody treatments.

Most leading anti-SARS-CoV-2 antibodies target the viral receptor-binding domain (RBD), which mediates binding to ACE2 receptor (*5*, *6*). We recently developed a deep mutational scanning method to map how all mutations to the RBD affect its function and recognition by antiviral antibodies (*7*, *8*). This method involves creating libraries of RBD mutants, expressing them on the surface of yeast, and using fluorescence-activated cell sorting and deep sequencing to quantify how each mutation affects RBD folding, ACE2 affinity, and antibody binding (Fig. S1A). Here we applied this method to map all RBD mutations that escape binding by recombinant forms of the two antibodies in Regeneron’s REGN-COV2 cocktail (REGN10933 and REGN10987) (*9*, *10*), and Eli Lilly’s LY-CoV016 antibody (also known as CB6 or JS016) (*11*) (Fig. S1B). REGN-COV2 was recently granted an emergency use authorization for treatment of COVID-19 (*12*), while LY-CoV016 is currently in phase 2 clinical trials (*13*).

We completely mapped RBD mutations that escape binding by the three individual antibodies as well as the REGN10933 + REGN10987 cocktail (Fig. 1A,B and zoomable maps at https://jbloomlab.github.io/SARS-CoV-2-RBD_MAP_clinical_Abs/). REGN10933 and REGN10987 are escaped by largely non-overlapping sets of mutations in the RBD’s receptor-binding motif (Fig. 1A), consistent with structural work showing that these antibodies target distinct epitopes in this motif (*9*). But surprisingly, one mutation (E406W) strongly escapes the cocktail of both antibodies (Fig. 1A). The escape map for LY-CoV016 also reveals escape mutations at a number of different sites in the RBD (Fig. 1B). Although some escape mutations impair the RBD’s ability to bind ACE2 or be expressed in properly folded form, many come at little or no cost to these functional properties (colors in Fig. 1A,B and Fig. S2)—an unfortunate consequence of the mutational tolerance of the RBD (*7*).

**Figure 1.**
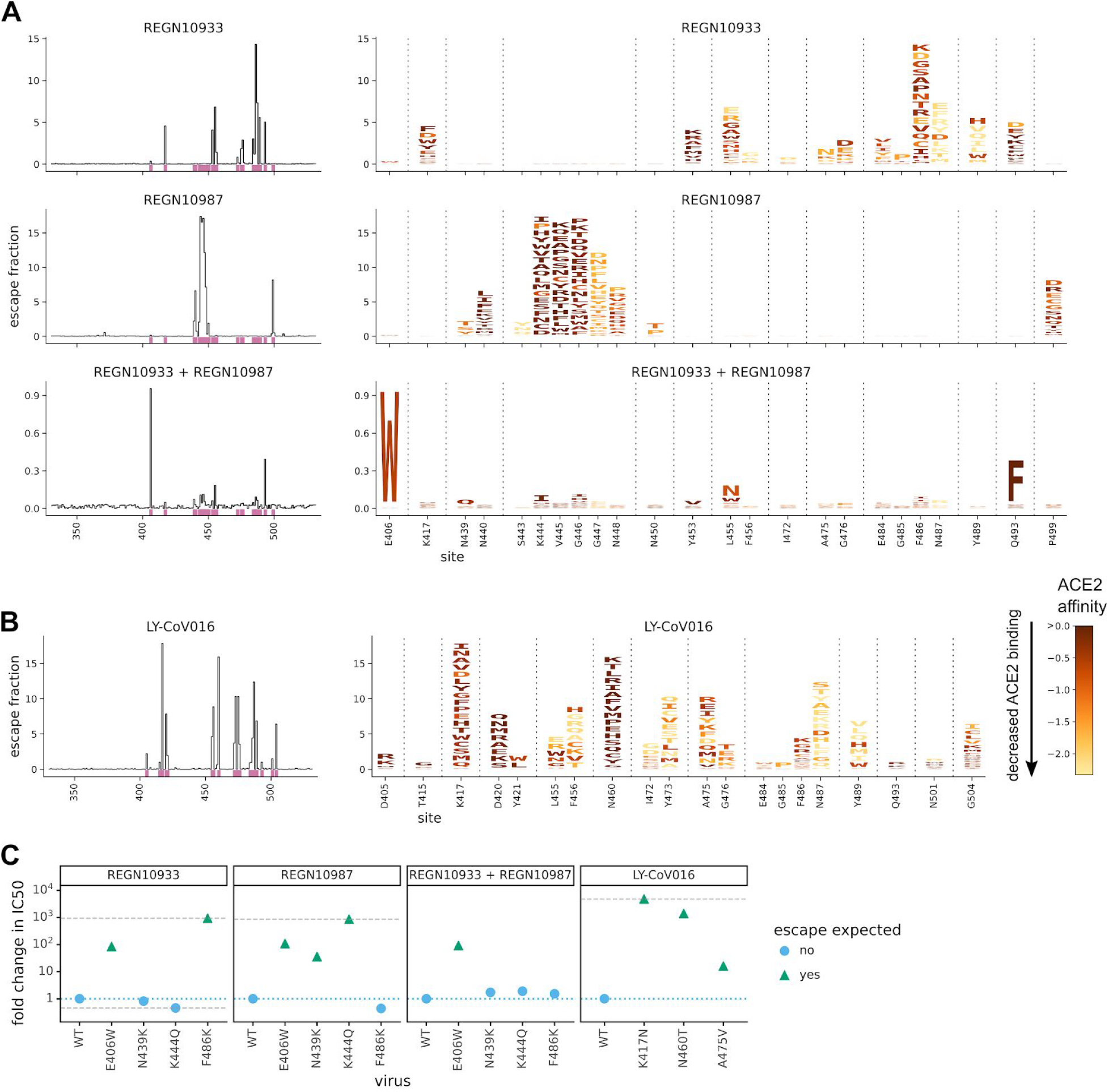
Complete maps of escape mutations from the REGN-COV2 antibodies and Ly-CoV016. (A) Maps for antibodies in REGN-COV2. Line plots at left show total escape at each site in the RBD. Sites of strong escape (purple underlines) are shown in logo plots at right. The height of each letter is proportional to how strongly that amino-acid mutation mediates escape, with a per-mutation “escape fraction” of 1 corresponding to complete escape. The y-axis scale is different for each row, so for instance E406W escapes all REGN antibodies but it is most visible for the cocktail as it is swamped out by other sites of escape for the individual antibodies. See https://jbloomlab.github.io/SARS-CoV-2-RBD_MAP_clinical_Abs/ for zoomable versions. Letters are colored by how mutations affect the RBD’s affinity for ACE2 (*7*), with yellow indicating poor affinity and brown indicating good affinity; see Fig. S2 for maps colored by how mutations affect expression of folded RBD. (B) Map for LY-CoV016. (C) Validation of key mutations in neutralization assays using pseudotyped lentiviral particles. Each point indicates the fold-increase in inhibitory concentration 50% (IC50) for a mutation relative to the unmutated “wildtype” (WT) Wuhan-Hu-1 RBD. The dotted blue line indicates wildtype-like neutralization sensitivity, and the dashed gray lines indicate upper and lower bounds on detectable fold changes. Point shapes / colors indicate if escape was expected at that site from the maps. Full neutralization curves are in Fig. S3.

To validate the antigenic effects of key mutations, we performed neutralization assays using spike-pseudotyped lentiviral particles, and found concordance between the escape maps and neutralization assays (Fig. 1C and Fig. S3). As expected from the maps for the REGN-COV2 antibodies, a mutation at site 486 escaped neutralization only by REGN10933, whereas mutations at sites 439 and 444 escaped neutralization only by REGN10987—and so none of these mutations escaped the cocktail. But E406W escaped both individual REGN-COV2 antibodies, and thus also strongly escaped the cocktail. The identification of E406W as a cocktail escape mutation demonstrates how complete maps provide information beyond other standard approaches: structural analyses and viral-escape selections led Regeneron to posit that no single amino-acid mutation could escape both antibodies in the cocktail (*9*, *10*), but our complete maps show this is not true.

To explore how well our escape maps explain the evolution of virus under antibody selection, we first examined data from Regeneron’s viral escape-selection experiments in which spike-expressing VSV was grown in cell culture in the presence of REGN10933, REGN10987, or the cocktail (*10*). That work identified five escape mutations from REGN10933, two from REGN10987, and none from the cocktail (Fig. 2A). All five cell-culture-selected mutations were prominent among the single-nucleotide accessible mutations in our escape maps (Fig. 2B), demonstrating concordance between the escape maps and viral evolution under antibody pressure in cell culture. Notably, E406W is not accessible by a single-nucleotide change, which may explain why it was not identified by the Regeneron cocktail selections despite being relatively well tolerated for RBD folding and ACE2 affinity.

**Figure 2.**
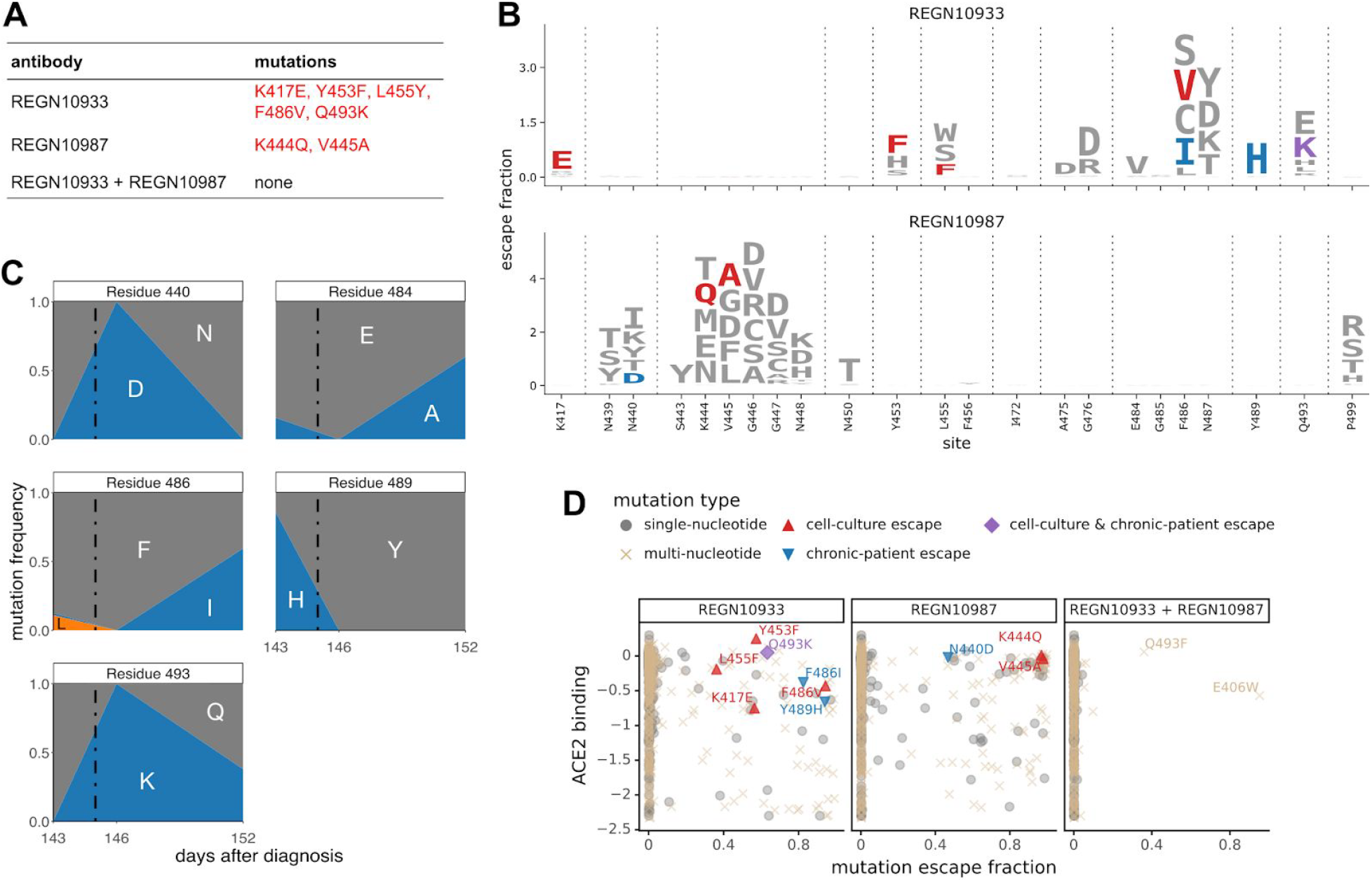
Escape maps are consistent with viral mutations selected in cell culture and a persistently infected patient. (A) Viral escape mutations selected by Regeneron with spike-pseudotyped VSV in cell culture in the presence of antibody (*10*). (B) Escape maps like those in Fig. 1A but showing only mutations accessible by single-nucleotide changes to the Wuhan-Hu-1 sequence, with non-gray colors indicating mutations in cell culture (red), in the infected patient (blue), or both (purple). Fig. S5 shows these maps colored by how mutations affect ACE2 affinity or RBD expression. (C) Dynamics of RBD mutations in a patient treated with REGN-COV2 at day 145 of his infection (black dashed vertical line). E484A rose in frequency in linkage with F486I, but since E484A is not an escape mutation in our maps it is not shown in other panels. See also Fig. S4. (D) The escape mutations that arise in cell culture and the infected patient are single-nucleotide accessible and escape antibody binding without imposing a large cost on ACE2 affinity. Each point is a mutation with shape / color indicating whether it is accessible and selected during viral growth. Points further to the right on the x-axis indicate stronger escape from antibody binding; points further up on the y-axis indicate higher ACE2 affinity.

To determine if the escape maps could also inform analysis of viral evolution in infected humans, we examined deep sequencing data from a persistently infected immunocompromised patient who was treated with REGN-COV2 at day 145 after diagnosis with COVID-19 (*14*). The late timing of treatment allowed ample time for the patient’s viral population to accumulate genetic diversity. Administration of REGN-COV2 was followed by rapid changes in the frequencies of five amino-acid mutations in the RBD (Fig. 2C and Fig. S4). Our escape maps showed that three of these mutations escaped REGN10933, and one escaped REGN10987 (Fig. 2B). Notably, the mutations did not all sweep to fixation after antibody treatment: instead, there were competing rises and falls (Fig. 2C). This pattern has been observed in the adaptive within-host evolution of other viruses (*15*, *16*), and occurs because of genetic hitchhiking and competition among viral lineages. Both these forces are apparent in the persistently infected patient (Fig. 2C and Fig S4C): E484A (not an escape mutation in our maps) hitchhikes with F486I (which escapes REGN10933) after treatment, and the viral lineage carrying N440D and Q493K (which escape REGN10987 and REGN10933, respectively) competes first with the REGN10933 escape-mutant Y489H, and then with the E484A / F486I lineage and Q493K-alone lineage.

Importantly, three of the four escape mutations in the REGN-COV2-treated patient were not identified in Regeneron’s viral cell-culture selections (Fig. 2B), illustrating an advantage of complete maps. Viral selections are “incomplete” in the sense that they only identify whatever mutations are stochastically selected in that particular cell-culture experiment. In contrast, complete maps annotate all mutations, which could include mutations that arise for reasons unrelated to treatment but incidentally affect antibody binding.

Of course, viral evolution is shaped by functional constraints as well as pressure to evade antibodies. The mutations selected in cell culture and the patient consistently met the following criteria: they escaped antibody binding, were accessible via a single-nucleotide change, and imposed little or no cost on ACE2 affinity (as measured by prior deep mutational scanning (*7*); Fig. 2D, Fig. S5). Therefore, complete maps of how mutations affect key biochemical phenotypes of the RBD (e.g., ACE affinity and antibody binding) can be used to assess likely paths of viral evolution. A caveat is that over longer evolutionary timeframes, the space of tolerated mutations could shift due to epistatic interactions, as has been previously observed in viral immune and drug escape (*17*–*19*).

The complete maps enable us to assess what escape mutations are already present among circulating SARS-CoV-2. We examined all human-derived SARS-CoV-2 sequences available as of November 12, 2020, and found a substantial number of RBD mutations that escaped one or more of the antibodies (Fig. 3). However, the only escape mutations present in >0.1% of sequences were the REGN10933 escape-mutant Y453F (0.2% of sequences) (*10*) and the REGN10987 escape-mutant N439K (1.2% of sequences, has an effect on neutralization as shown in both Fig. 1C and (*20*)). Y453F is associated with independent mink-associated outbreaks in the Netherlands and Denmark (*22*, *23*); notably the mink sequences themselves sometimes also contain other escape mutations such as F486L (*21*). N439K is prevalent in Europe, where it has comprised a large percentage of sequences from regions including Scotland and Ireland (*20*, *23*).

**Figure 3.**
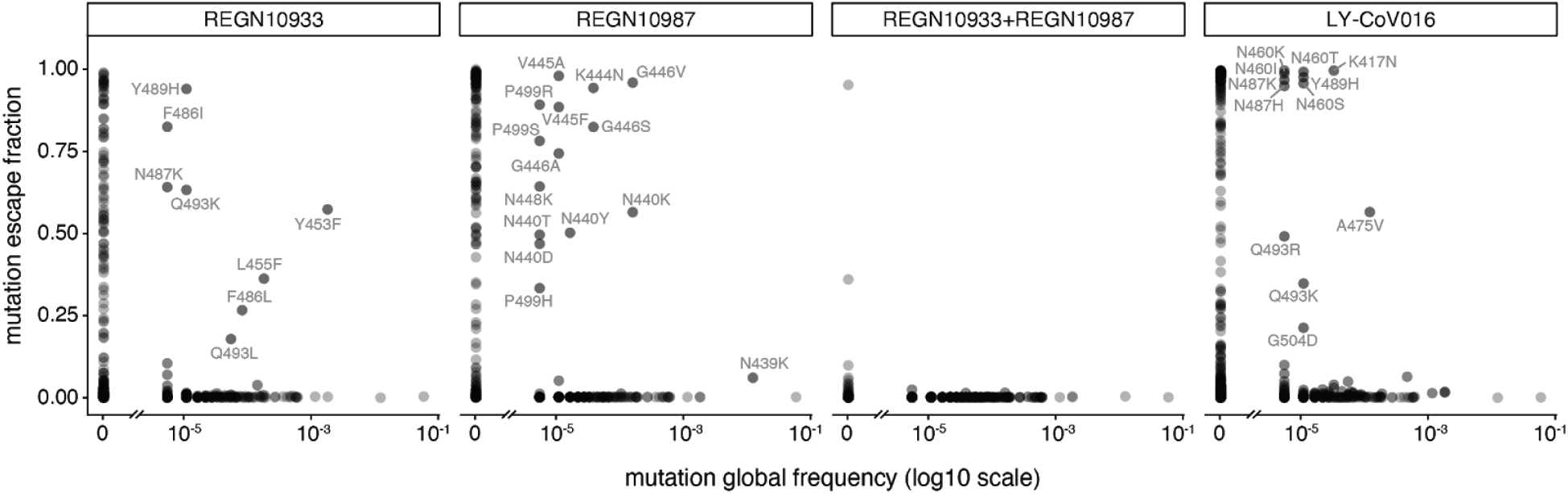
Antibody escape mutations in circulating SARS-CoV-2. For each antibody or antibody combination, the escape score for each mutation is plotted versus its frequency among the 180,555 high-quality human-derived SARS-CoV-2 sequences on GISAID (*25*) as of November 12, 2020. Escape mutations with notable GISAID frequencies are labeled.

To determine if the escape maps could be rationalized from the structural interfaces of the antibodies and RBD, we projected the maps onto crystal or cryo-EM structures (Fig. 4A; interactive versions at https://jbloomlab.github.io/SARS-CoV-2-RBD_MAP_clinical_Abs/). As might be expected, escape mutations generally occur in the antibody-RBD interface. However, structures alone are insufficient to predict which mutations mediate escape. For example, LY-CoV016 uses both its heavy and light chains to bind a wide epitope overlapping the ACE2-binding surface, but escape is dominated by mutations at RBD residues that contact the heavy chain CDRs (Figs. 4A, S6E-G). In contrast, escape from REGN10933 and REGN10987 mostly occurs at RBD residues that pack at the antibody heavy/light-chain interface (Fig. 4A, S6A-D). The E406W mutation that escapes the REGN-COV2 cocktail occurs at a residue not in contact with either antibody (Fig. 4A). So overall, mutations at RBD residues that contact antibody do not always mediate escape, and several prominent escape mutations occur at residues not in contact with antibody (Fig. 4B, S6D,G).

**Figure 4.**
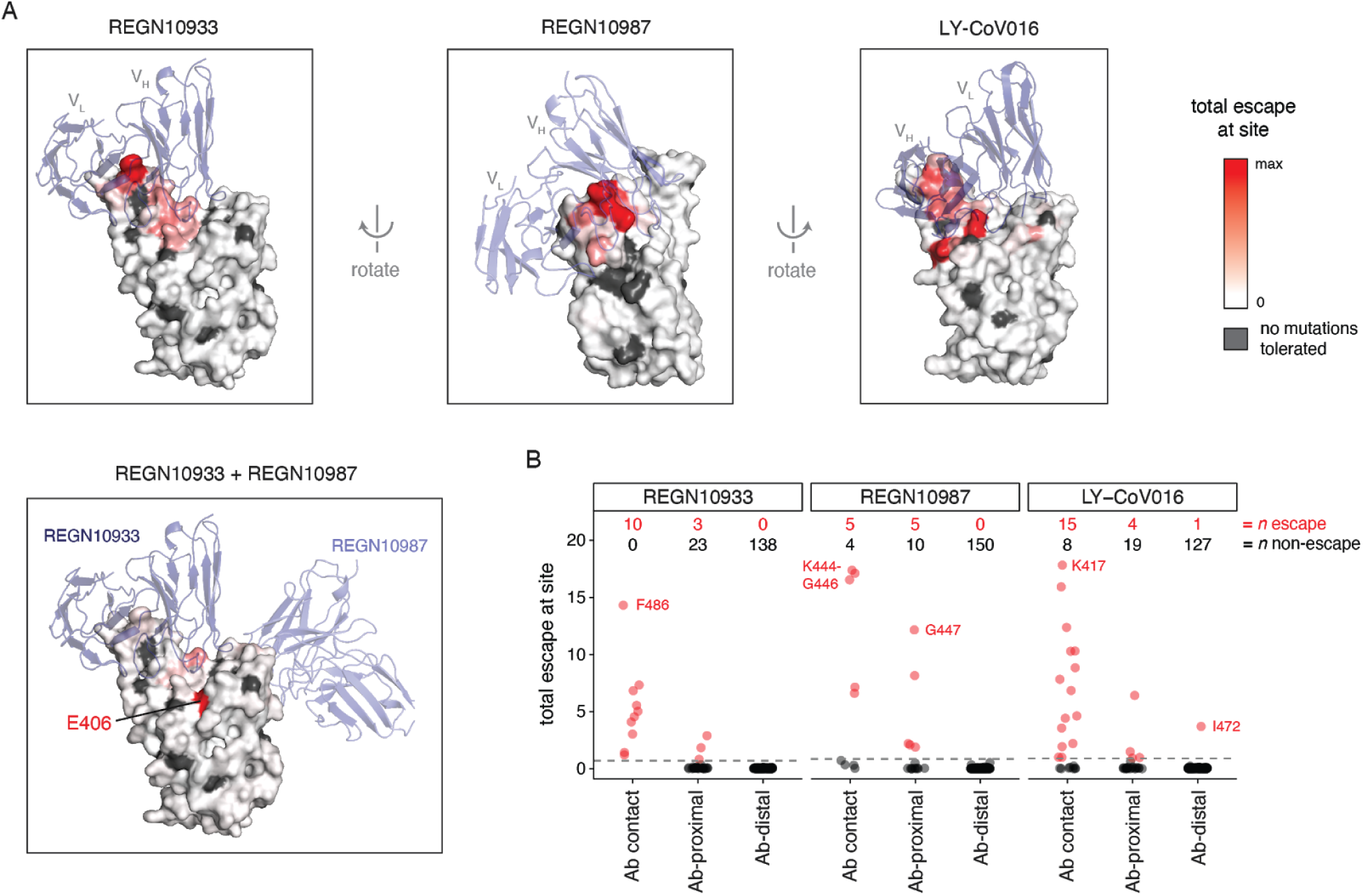
Structural context of escape mutations. (A) Escape maps projected on antibody-bound RBD structures. (REGN10933 and REGN10987: PDB 6XDG (*9*); LY-CoV016: PDB 7C01 (*11*)). Antibody heavy- and light-chain variable domains are shown as blue cartoons, and the RBD surface is colored to indicate how strongly mutations at that site mediate escape (white indicates no escape, red indicates strongest escape site for that antibody / cocktail). Sites where no mutations are functionally tolerated are colored gray. (B) For each antibody, sites were classified as direct antibody contacts (non-hydrogen atoms within 4Å of antibody), antibody-proximal (4-8Å), or antibody-distal (>8Å). Each point indicates a site, classified as escape (red) or non-escape (black) (dashed gray line, see Methods). Red and black numbers indicate how many sites in each category are escape or non-escape, respectively. Interactive visualizations are at https://jbloomlab.github.io/SARS-CoV-2-RBD_MAP_clinical_Abs/ and additional static views are in Fig. S6.

Overall, we have completely mapped mutations that escape some of the leading antibodies used to treat COVID-19. These maps demonstrate that prior characterization of escape mutations was incomplete: for instance, overlooking a single amino-acid mutation that escapes both antibodies in the REGN-COV2 cocktail, and failing to identify most mutations that arose in a persistently infected patient treated with the cocktail. Of course, our maps still do not answer the most pressing question: will SARS-CoV-2 evolve widespread resistance to these antibodies? While the presence of escape mutations in the patient treated with REGN-COV2 is ominous, other viruses that typically cause self-limiting acute infections undergo extensive within-patient evolution only in long infections of immunocompromised patients (*15*) and not in the broader population (*24*). However, it is concerning that so many escape mutations impose little cost on RBD folding or receptor affinity, and that some of these mutations are already present at low levels among circulating viruses. Ultimately, it will be necessary to wait and see what mutations spread as SARS-CoV-2 circulates in the human population. Our work will help with the “seeing,” by enabling immediate interpretation of the effects of the mutations catalogued by viral genomic surveillance.

## Acknowledgements

We thank Katharine Crawford for help with neutralization assays, Alison Feder for helpful comments, and the Fred Hutch Flow Cytometry and Genomics facilities for assistance.

## Author contributions

TNS, AJG, ASD, and JDB designed the study. TNS, AJG, AA, and ASD performed the experiments. TNS, AJG, and JDB analyzed the experimental data. JZL and MCC sequenced the persistent infection, and WWH analyzed that data. TNS, AA, WWH, and JDB wrote the initial draft, and all authors edited the final version.

## Data and materials availability

Raw sequencing data are on the NCBI SRA under BioProject PRJNA639956 / BioSample SAMN16850904 (escape mapping) and Bioproject PRJNA681234 (patient sequencing). Computer code and processed data are on GitHub: https://github.com/jbloomlab/SARS-CoV-2-RBD_MAP_clinical_Abs (escape mapping) and https://github.com/jbloomlab/SARS-CoV-2_chronic-infection-seq (patient sequencing). See Materials and Methods for links to specific analyses.

## Supplementary Materials

## Materials and Methods

### Data and Code Availability

- Complete computational pipeline for escape-mapping data analysis: https://github.com/jbloomlab/SARS-CoV-2-RBD_MAP_clinical_Abs
- Markdown summaries of the escape-mapping data analysis steps: https://github.com/jbloomlab/SARS-CoV-2-RBD_MAP_clinical_Abs/blob/main/results/summary/summary.md
- Raw data tables of mutant escape fractions: https://github.com/jbloomlab/SARS-CoV-2-RBD_MAP_clinical_Abs/blob/main/results/supp_data/REGN_and_LY-CoV016_raw_data.csv
- Raw Illumina sequencing for the escape mapping: NCBI SRA, BioProject: PRJNA639956, BioSample SAMN16850904
- Processed Illumina sequencing counts for the escape mapping: https://github.com/jbloomlab/SARS-CoV-2-RBD_MAP_clinical_Abs/tree/main/results/counts
- Complete computational pipeline for analysis of within-patient viral evolution: https://github.com/jbloomlab/SARS-CoV-2_chronic-infection-seq
- Raw Illumina sequencing for the within-patient viral evolution: NCBI SRA, BioProject PRJNA681234.

### Antibodies

Publicly available antibody variable domain sequences were acquired for REGN10933, REGN10987, and LY-CoV016 (also known as JS016, LY3832479, or CB6). Specifically, REGN10933 and REGN10987 variable domain sequences were reported by Hansen et al. (*9*) in supplemental Data S1. LY-CoV016 (CB6) sequence was reported by Shi et al. (*11*), Genbank Accessions MT470196 and MT470197.

Recombinant antibodies were cloned and produced by Genscript. Specifically, antibody variable domains were cloned with the human IgG1 heavy chain and human IgK (REGN10933 and LY-CoV016) or human IgL2 (REGN10987) constant regions into pcDNA3.4 vector, and transfected into HD 293F cells maintained at 37°C with 8% CO2 on an orbital shaker. Cell culture supernatants were collected, and affinity purified over RoboColumn Eshmuno A 0.6mL columns.

### Antibody-escape mapping

Antibody selection experiments were performed in biological duplicate using a deep mutational scanning (mutational antigenic profiling) approach (*8*) using previously described duplicate mutant RBD libraries (*7*). These libraries contain virtually all possible amino-acid mutations to the SARS-CoV-2 RBD within a yeast-surface display vector, with RBD variants linked to unique 16-nucleotide barcode sequences to facilitate downstream sequences. As described in (*8*), these libraries were sorted to eliminate variants that lose ACE2 binding prior to mapping the antibody-escape variants.

Antibody labeling and selection was performed essentially as described in (*8*). Specifically, 9 OD aliquots of RBD libraries were thawed and grown overnight at 30°C 275 rpm in 45mL SD-CAA (6.7 g/L Yeast Nitrogen Base, 5.0 g/L Casamino acids, 1.065 g/L MES, and 2% w/v dextrose). Libraries were backdiluted to an OD of 0.67 in SG-CAA+0.1% dextrose (SD-CAA with 2% w/v galactose and 0.1% w/v dextrose in place of 2% dextrose), and incubated for 16-18 hours at room temperature with mild agitation to induce RBD surface expression. For each antibody selection, 20 OD units of induced cells were washed twice with PBS-BSA (0.2 mg/mL), and incubated in 4mL PBS-BSA with 400 ng/mL antibody (monoclonal REGN10933, REGN10987, LY-CoV016, or REGN10933+REGN10987 pooled at 1:1 w/w ratio at total 400 ng/mL) for 1 h at room temperature with gentle agitation. Labeled cells were washed with ice-cold PBS-BSA followed by secondary labeling for 1 h at 4°C in 2.5 mL 1:200 PE-conjugated goat anti-human-IgG (Jackson ImmunoResearch 109-115-098) to label for bound antibody, and 1:100 FITC-conjugated anti-Myc (Immunology Consultants Lab, CYMC-45F) to label for RBD surface expression. Labeled cells were washed twice with PBS-BSA and resuspended in 2.5mL PBS. Yeast expressing the unmutated SARS-CoV-2 RBD were prepared in parallel to library samples, labeled at the same 400 ng/mL and 100x reduced 4 ng/mL antibody concentrations.

Antibody-escape cells were selected via fluorescence-activated cell sorting (FACS) on a BD FACSAria II. FACS selection gates were drawn to capture 95% of yeast expressing unmutated SARS-CoV-2 RBD labeled at 4 ng/mL antibody (100x reduced antibody concentration relative to library samples, see Figure S1B,C). For each library sample, approximately 6-8 million RBD+ cells were processed on the cytometer, with between 5.9e5 and 1.9e6 antibody-escaped cells collected per sample into SD-CAA supplemented with 1% w/v BSA (see selection percentages in Figure S1C). Antibody-escaped cells were grown overnight in 1.5mL SD-CAA + 100 U/mL penicillin + 100 μg/mL streptomycin at 30°C 275 rpm.

Plasmid samples were prepared from up to 7.5 OD units of overnight cultures of antibody-escaped cells, and 30 OD units of pre-selection yeast populations (Zymoprep Yeast Plasmid Miniprep II) per manufacturer instructions, with the addition of a −80°C freeze-thaw step prior to cell lysis. The 16-nucleotide barcode sequences identifying each RBD variant were amplified by PCR and prepared for Illumina sequencing exactly as described by Starr et al. (*7*). Barcodes were sequenced via 50 bp single-end reads on an Illumina HiSeq 3500, targeting at least 3x as many sequencing reads as FACS-selected cells, and pre-sort reference populations of at least 2.5e7 reads.

### Analysis of mutant library deep sequencing and computation of per-mutant escape fractions

Escape fractions were computed as described in (*8*), with minor modifications as noted below. Specifically, we used the dms_variants package (https://jbloomlab.github.io/dms_variants/, version 0.8.2) to process Illumina sequences into counts of each barcoded RBD variant in each pre-sort and antibody-escape population using the barcode/RBD look-up table from (*7*). Markdown renderings of these steps in the computational analysis are at https://github.com/jbloomlab/SARS-CoV-2-RBD_MAP_clinical_Abs/blob/main/results/summary/aggregate_variant_counts.md and https://github.com/jbloomlab/SARS-CoV-2-RBD_MAP_clinical_Abs/blob/main/results/summary/counts_to_cells_ratio.md.

For each antibody selection, we then computed the “escape fraction” for each barcoded variant using the deep sequencing counts for each variant in the original and antibody-escape populations and the total fraction of the library that escaped antibody binding via the formula provided in Greaney et al (*8*). These escape fractions represent the estimated fraction of cells expressing that specific variant that fall in the antibody escape bin, so a value of 0 means the variant is always bound by antibody and a value of 1 means that it always escapes antibody binding. We then applied a computational filter to remove variants with low sequencing counts or highly deleterious mutations that might cause antibody escape simply by leading to poor expression of properly folded RBD on the yeast cell surface. Specifically, we ignored all variants with pre-selection sequencing counts that were lower than the counts for the 99th percentile of the stop-codon containing variants--the logic here being that stop codon variants are largely purged by the earlier sorts for RBD expressing and ACE2-binding variants and so any residual presence provides an indication of low-count “noise.” Next, we removed any variants that had poor RBD expression or ACE2 binding, or contained mutations that individually cause poor RBD expression and ACE2 binding, the logic being that this would eliminate misfolded or non-expressing RBDs. Specifically, we removed variants that had (or contained mutations with) ACE2 binding scores < −2.35 or expression scores < −1, using the variant- and mutation-level deep mutational scanning scores from Starr et al (*8*). Note that these filtering criteria are slightly more stringent than those used in Greaney et al (*8*). A markdown rendering of the computation of the variant-level escape fractions and the variant filtering is at https://github.com/jbloomlab/SARS-CoV-2-RBD_MAP_clinical_Abs/blob/main/results/summary/counts_to_scores.md.

We next deconvolved variant-level escape scores into escape fraction estimates for single mutations using global epistasis models (*26*) implemented in the dms_variants package, as detailed at (https://jbloomlab.github.io/dms_variants/dms_variants.globalepistasis.html). In this fitting, we excluded variants that contained mutations that were not seen as either single mutants or in at least two multiple-mutant variants. We then computed the estimated effect of each mutation as the impact of that mutation on the “observed phenotype” scale transformation of its “latent phenotype” as computed using the global epistasis models, and applied a floor of zero and a ceiling of 1 to these escape fractions. All of the above analysis steps were performed separately for each of the duplicate mutant libraries. We then only retained those mutations that passed all of the above filtering and were measured in both libraries or had at least two-single mutant measurements in one library. The reported scores throughout the paper are the average across the libraries; these scores are also in Table S1. Correlations in final single-mutant escape scores are shown in Figure S1D. A markdown rendering of the computation that computes these mutation-level escape fractions is at https://github.com/jbloomlab/SARS-CoV-2-RBD_MAP_clinical_Abs/blob/main/results/summary/scores_to_frac_escape.md.

For plotting and analyses that required identifying RBD sites of “strong escape” (e.g., choosing which sites to show in logo plots in Fig 1A,B or label in Figure 4B), we considered a site to mediate strong escape if the total escape (sum of mutation-level escape fractions) for that site exceeded the median across sites by >5 fold, and was at least 5% of the maximum for any site. A markdown rendering of the identification of these sites of strong escape is at https://github.com/jbloomlab/SARS-CoV-2-RBD_MAP_clinical_Abs/blob/main/results/summary/call_strong_escape_sites.md.

### Pseudotyped lentiviral particle neutralization assays

We performed neutralization assays using lentiviral particles carrying the luciferase gene and pseudotyped with the SARS-CoV-2 spike essentially as described in Crawford et al (*27*) with the following two modifications: the Wuhan-Hu-1 spike sequence had a deletion of the final 21 amino acids in the cytoplasmic tail (which increases viral titers (*28*)), and carried the D614G mutation (which further increases viral titers and makes the sequence better match currently circulating viruses (*8*)). The spike plasmid used for these experiments, HDM-SARS2-spike-del21-D614G, is available on AddGene as plasmid #158762 (https://www.addgene.org/158762/).

### Deep-sequencing analysis of within-host viral genetic diversity in persistently infected patient

The persistently infected patient and his clinical time course are described in detail in (*14*). That paper also describes the Illumina deep sequencing of that patient at nine timepoints. All sequencing is from nasal swab samples. The deep sequencing data have been deposited on the Sequence Read Archive under BioProject accession PRJNA681234.

Intra-patient single-nucleotide polymorphisms (SNPs) were identified with an automated variant-calling pipeline (https://github.com/jbloomlab/SARS-CoV-2_chronic-infection-seq) created with Snakemake (*29*). Briefly, paired-end reads were filtered, and sequencing adaptors were removed with fastp (*30*). Reads from SARS-CoV-2 were enriched by kmer matching to the Wuhan-Hu-1 reference genome (NC_045512.2) using BBDuk (https://jgi.doe.gov/data-and-tools/bbtools/bb-tools-user-guide/). Following filtering, reads were aligned to the Wuhan-Hu-1 reference with BWA-MEM (*31*). Variants were identified by counting the coverage of each base at every position in the reference genome using a custom Python script. These variants were filtered based on a minimum allele frequency of >0.01, a PHRED quality threshold of >25, and coverage of more than 100 reads. The coverage pattern over the Spike gene was plotted by averaging the number of reads over every base meeting the minimum PHRED score of 25 in 10 bp bins (Fig. S4A).

To visualize the change in allele frequencies over time (Fig. 2C & S4B), we identified sites in the spike gene with nonsynonymous mutations that rose above 10% frequency at any sampled timepoint (note that we ignore any mutations relative to Wuhan-Hu-1 that are fixed at all timepoints as these are not intra-host variants). Using this list of high-confidence polymorphisms, we selected any other nonsynonymous mutations annotated at those sites, regardless of frequency, to get a full picture of allelic variation in putatively selected residues. For the analysis of just the RBD mutations between days 143 and 152 (Fig. 2C), we excluded any mutations that were either fixed or absent over the timeframe of interest (T478K, S494P, and N501Y).

To phase the variant alleles in the RBD (Fig. S4C) over the last three timepoints, we used a custom Python script that counted the co-occurrence of nonsynonymous variants in read-pairs. To maximize the number of informative reads for each timepoint, we only required that reads cover segregating sites in each timepoint based on analysis of the mutation frequencies in Fig. 2C. In other words, for the day 143 sample, we required reads to cover sites 484, 486, and 489, but not sites 440 or 493. For the day 146 sample, only one haplotype was possible (N440D/Q493K); thus, its frequency was assumed to be 100%. Finally, for the day 152 sample, we required reads to cover sites 484, 486, 489, and 493, but not site 440. Of these informative reads, those with SAM flags indicating quality failure or secondary mapping were excluded. To estimate the frequency of the identified haplotypes, we divided each haplotype’s count by the total number of unique haplotypes at each timepoint. Despite the lower number of supporting reads for each haplotype than for individual variants (527 reads for Day 143; 732 reads for Day 146; 1091 reads for Day 152), each haplotype’s frequencies were consistent with the frequencies of the individual variants of which they were comprised. Finally, we filtered out any haplotypes present at a frequency of less than 0.01.

### Analysis of mutations in circulating human SARS-CoV-2 strains

For the analysis in Fig. 3, all 196,061 spike sequences on GISAID (*25*) as of 12-November-2020 were downloaded and aligned via mafft (*32*). Sequences from non-human origins and sequences containing gap or ambiguous characters were removed, as were sequences with extremely high numbers of RBD mutations relative to other sequences, leaving 180,555 retained sequences. All RBD amino-acid mutations were enumerated compared to the reference Wuhan-Hu-1 SARS-CoV-2 RBD sequence (Genbank MN908947, residues N331-T531). To explore the prevalence of mutations such as Y453F and N439K with finer-scale geographic resolution, we used the COVID-19 CG resource (covidcg.org) (*23*). We acknowledge all contributors to the GISAID EpiCoV database for their sharing of sequence data (all contributors listed at: https://github.com/jbloomlab/SARS-CoV-2-RBD_MAP_clinical_Abs/blob/main/data/gisaid_hcov-19_acknowledgement_table_2020_11_12.pdf).

### Data visualization

The static logo plots in the paper were created using `dmslogo` (https://jbloomlab.github.io/dmslogo/) version 0.5.0; a markdown rendering of the code that creates these logo plots is at https://github.com/jbloomlab/SARS-CoV-2-RBD_MAP_clinical_Abs/blob/main/results/summary/escape_profiles.md.

The interactive visualizations of the escape maps and their projections on the RBD-antibody structures available at https://jbloomlab.github.io/SARS-CoV-2-RBD_MAP_clinical_Abs/ were created using `dms-view` (https://dms-view.github.io/docs/) (*33*).

The static structural views in the paper were rendered in PyMOL using antibody-bound RBD structures PDB 6XDG (*9*) and PDB 7C01 (*11*). Structural distances were computed using the bio3d package in R (*34*).

---

**Figure S1.**
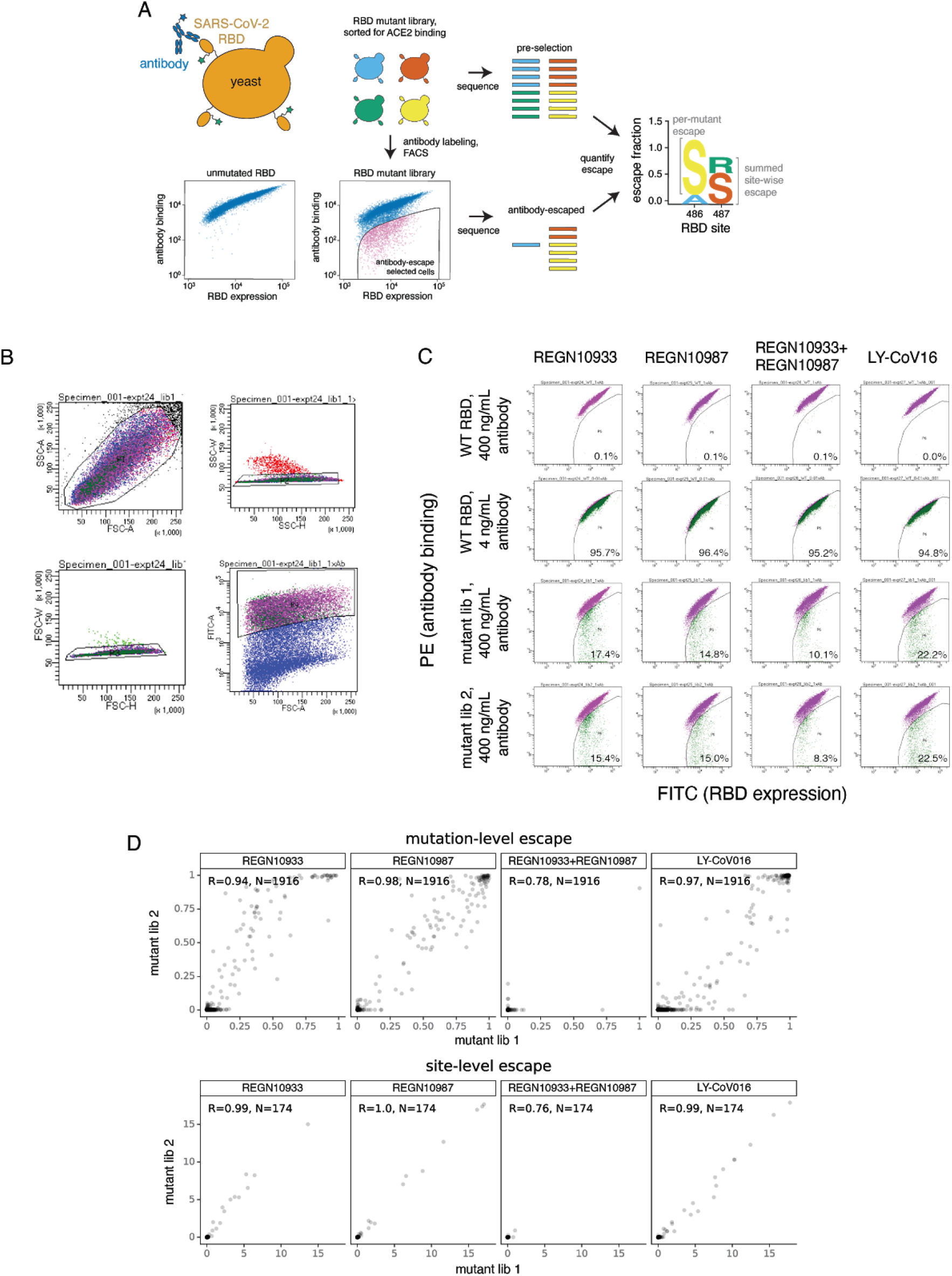
Deep mutational scanning method to map antibody-escape mutations. (A) Experimental approach to map antibody-escape mutations (*8*). SARS-CoV-2 RBD is expressed on the yeast cell surface (*7*), where fluorescent labeling detects RBD surface expression and antibody binding. A library of the SARS-CoV-2 RBD variants, previously sorted to purge non-functional variants (*8*), is labeled with antibody. Individual yeast cells expressing antibody-escape RBD variants are isolated via fluorescence-activated cell sorting (FACS). Deep sequencing quantifies variant frequencies before and after FACS, enabling the calculation of an “escape fraction” for each RBD mutation, which describes the fraction of cells containing a mutation that fall into the antibody-escape FACS bin. Escape fractions are illustrated in logoplots, where the height of a letter indicates the escape fraction for an individual mutation, and the sum of letter heights at a position indicates the total escape at a site. (B) Representative FACS gates used to select single yeast cells (nested SSC/FSC, SSC-W/SSC-H, and FSC-W/FSC-H gates) that express RBD on the cell surface (FITC/FSC). (C) Among RBD+ cells, antibody-escape bins were drawn on antibody-binding versus RBD expression scatterplots, with gate stringency determined from unmutated RBD controls. Antibody-escape sort gates were drawn to capture ~95% of cells expressing unmutated SARS-CoV-2 RBD when labeled at 0.01x the concentration of antibody used to label mutant libraries. The percentage of cells that fall in the antibody-escape bin in controls and independent library replicates are shown. (D) Correlations in deep mutational scanning scores between independent library duplicates. For each antibody, the escape fraction of individual mutations (top) and total escape per site (bottom) is shown for two independently generated and assayed mutant libraries. R, Pearson correlation coefficient. N, number of mutations or sites. Virtually all of the 3,819 possible RBD mutations are present in our libraries, but mutations that completely disrupt folding or binding are purged prior to antibody selections (see Methods).

**Figure S2.**
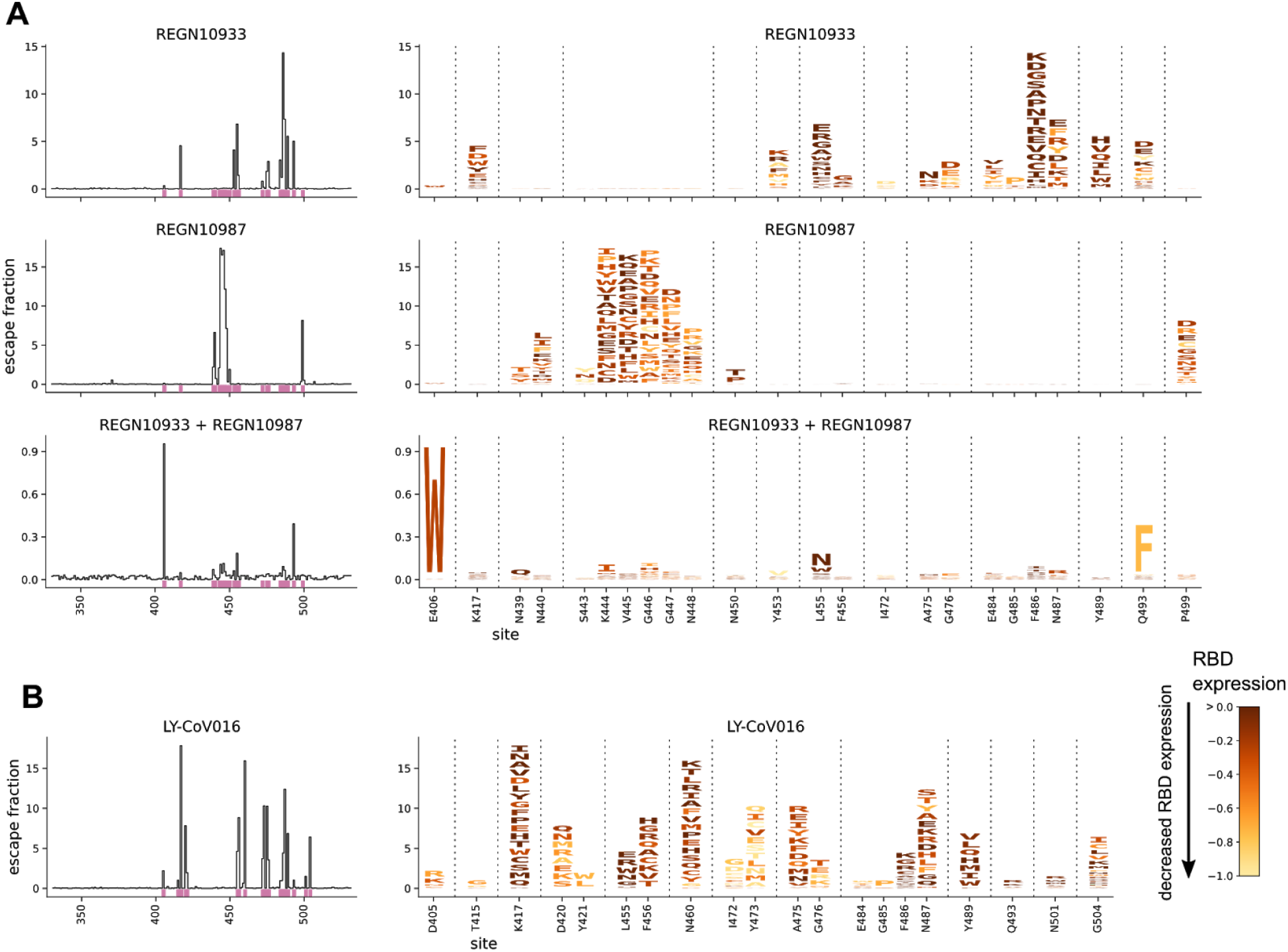
Complete escape maps colored by effects of mutations on RBD expression. The escape maps shown here are identical to those in Fig. 1A,B except that the letters are colored by how mutations affect RBD expression (*7*) rather than how they affect the RBD’s affinity for ACE2.

**Figure S3.**
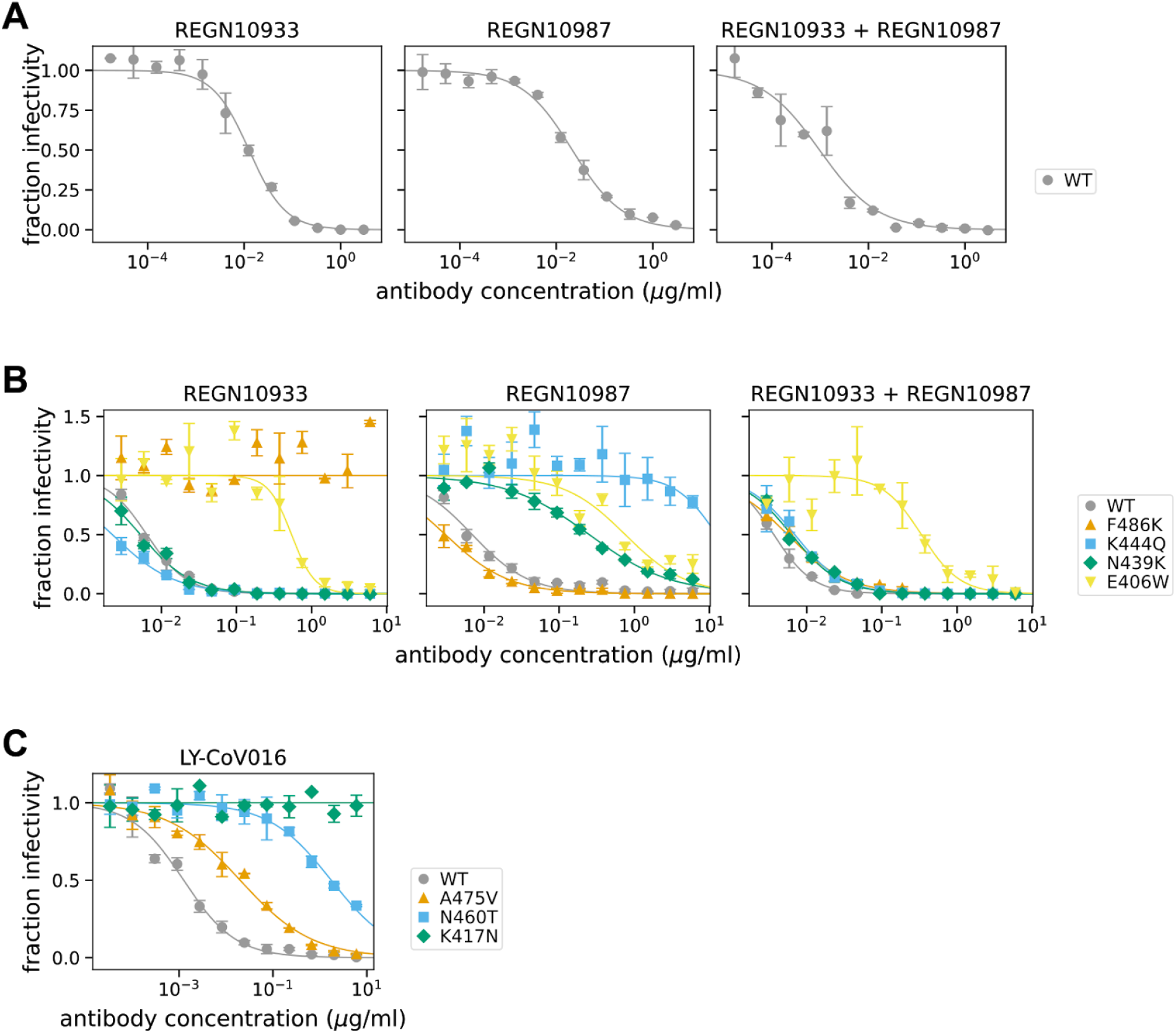
Pseudovirus neutralization curves validating the escape mutant mapping. (A) We first performed neutralization assays against the REGN-COV2 antibodies / cocktail using the unmutated SARS-CoV-2 spike to identify an appropriate dilution range to capture the inhibitory concentration 50% (IC50). (B) We then performed neutralization assays using the REGN-COV2 antibodies / cocktail with the indicated spike mutants, using a dilution range that spanned higher antibody concentration ranges to maximize the resolution on changes in IC50 for escape mutations. The changes in IC50 caused by the mutations as determined from these curves are what is shown in Fig. 1C. For the REGN10933 + REGN10987 cocktail, the concentration on the x-axis represents the total concentration of antibody, with the two components at an equimolar ratio. (C) Neutralization curves for LY-CoV016 against some of its key escape mutations.

**Figure S4.**
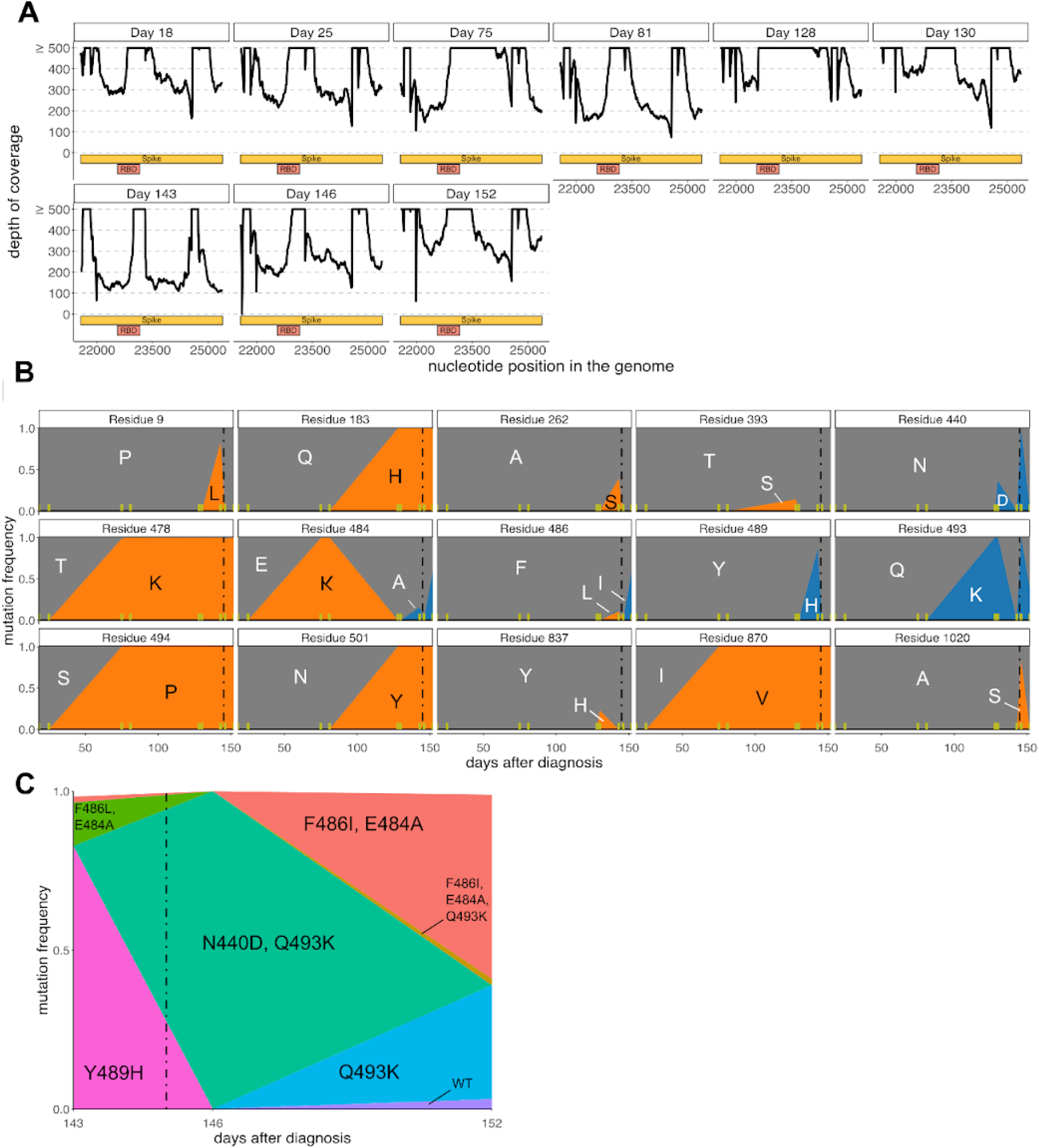
Spike mutations in a persistently infected patient treated with REGN-COV2 as determined by Illumina deep sequencing. **(A)** Coverage at each site in spike for each timepoint, calculated as the average number of aligned reads with a Q-score ≥25 in 10 bp bins. The x-axis shows the nt position in the genome coordinates of Wuhan-Hu-1 (NC_045512.2). The bars underneath show spike and its RBD domain. Coverage >500 is clipped on the y-axis. (B) Dynamics of amino-acid mutations in spike across all timepoints. Yellow vertical lines on the x-axis indicate sampling times, and the dashed black line indicates administration of REGN-COV2 (145 days). Fig. 1C is a subset of this plot that just shows RBD mutations in the timepoint immediately before and then after REGN-COV2 administration; those mutations are indicated in blue while all others are in orange. (C) Frequencies of different haplotypes in the RBD at the last three timepoints show competition among viral lineages. Note that it is possible that rare haplotypes (such as F468I / E484A / Q493K haplotype) represent library preparation artifacts that arise due to PCR strand exchange between molecules from more common haplotypes (e.g., F486I / E484A haplotype and Q493K haplotype).

**Figure S5.**
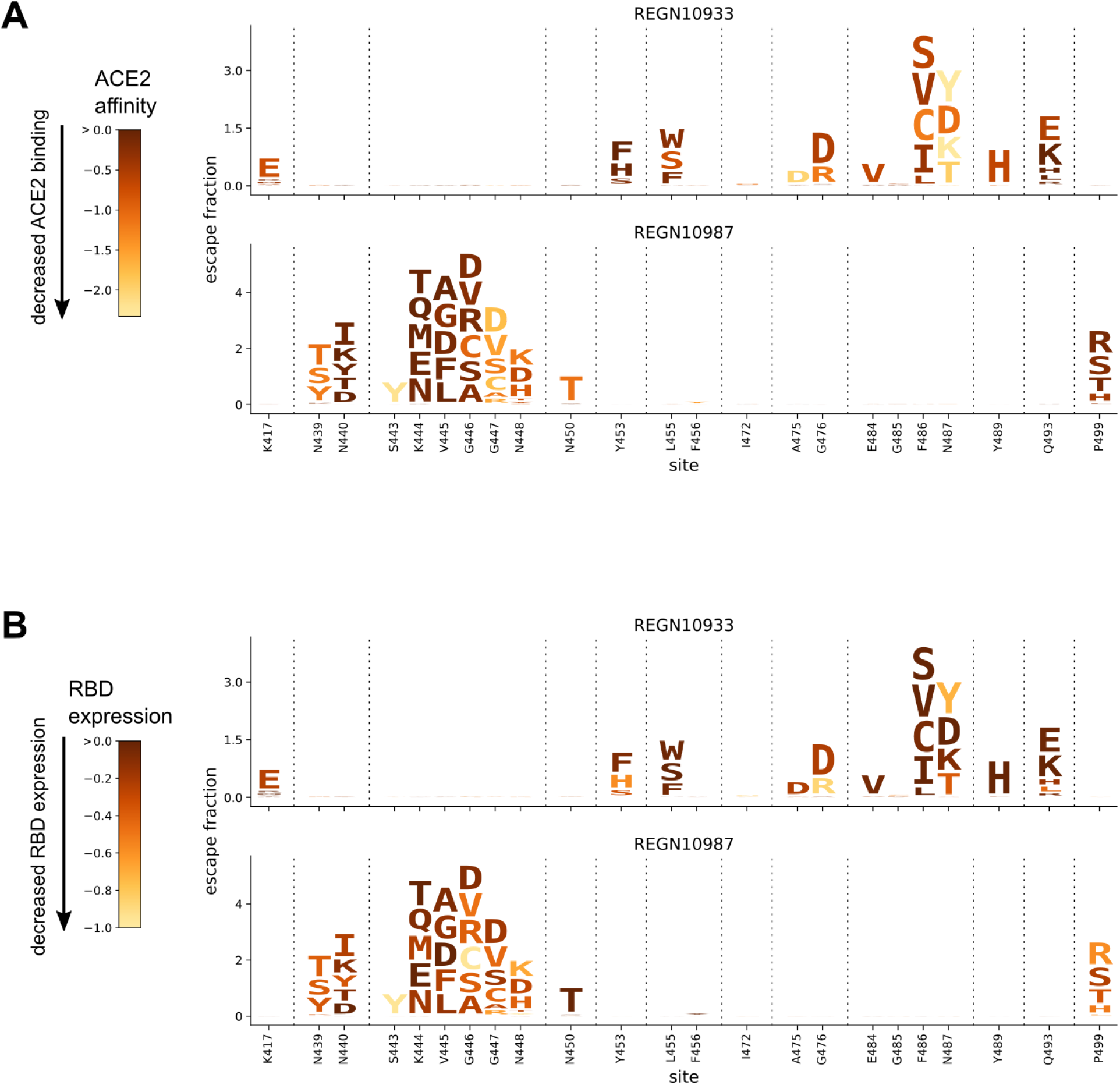
Maps of single-nucleotide accessible escape mutations from REGN10933 and REGN10987 colored by how mutations affect the RBD’s affinity for ACE2 or expression of folded protein. These plots show the same mutations as in Figure 2B (only those accessible by single-nucleotide changes to Wuhan-Hu-1), but colored according to the schemes in Fig 1A and Fig S2.

**Figure S6.**
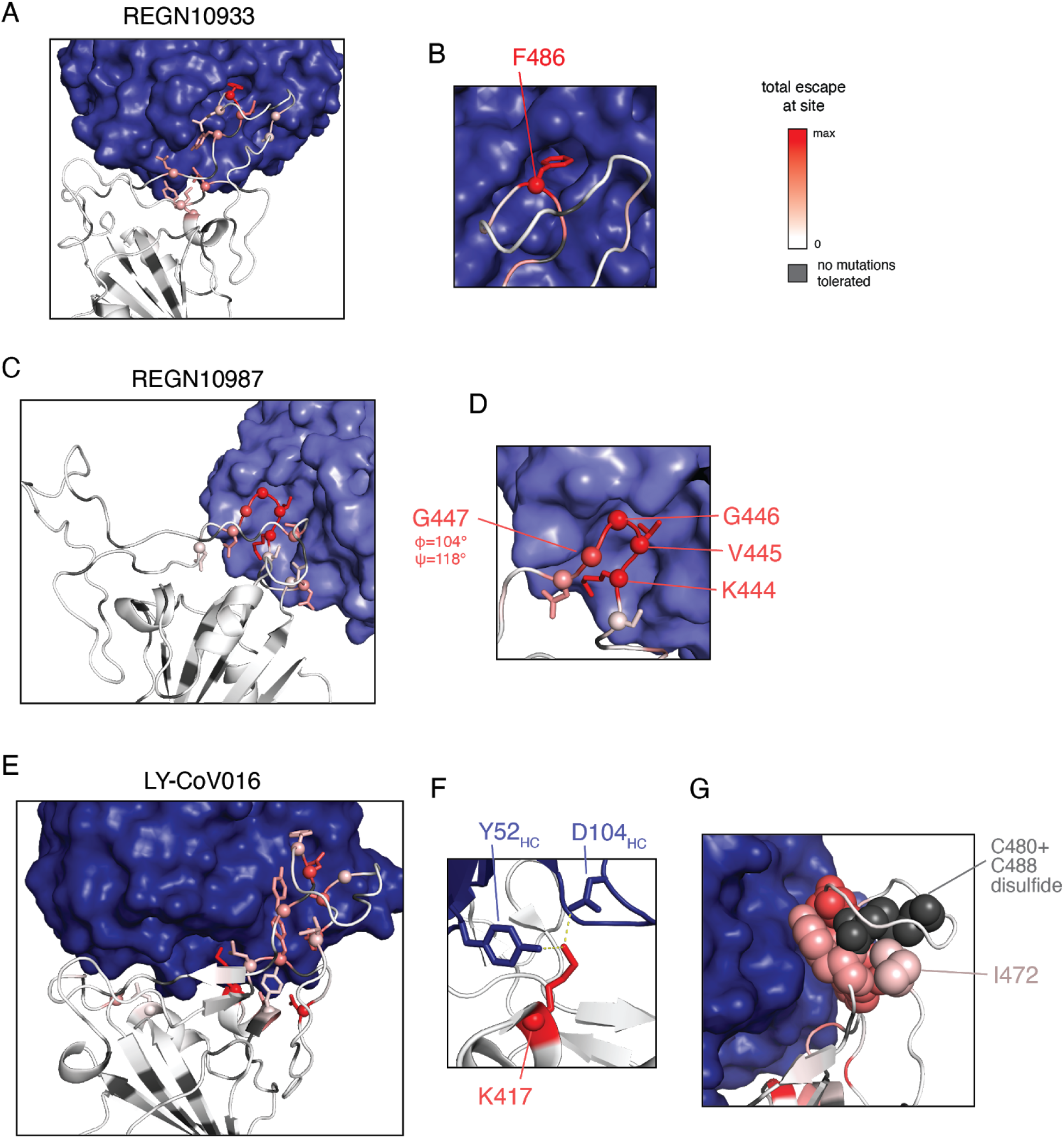
Structural mechanisms of escape. (A, C, E) Mapping of escape to antibody-bound RBD structures (PDB 6XDG (*9*), 7C01 (*11*)), with antibodies as blue surface and RBD colored by escape from white to red (see scale bar, upper right). RBD sites of escape are shown as sidechain sticks with spheres at alpha carbons. Zoomed views of sites of interest are presented to the right of each antibody structure. (B) F486, the top escape site for REGN10933, inserts into a large hydrophobic pocket at the antibody surface. (D) Residues K444, V445, and G446, the top sites of escape for REGN10987, are part of a loop that packs tightly with REGN10987. G447, a prominent site of escape that is not a direct contact (and mutant side chains point away from the antibody surface), is at the base of this loop and occupies a glycine-specific phi/psi conformational state. Mutations to G447 likely disturb the precise conformation of the 444-446 loop, thereby escaping REGN10987 binding. (F) K417, the top site of escape for LY-CoV016, forms polar contacts with antibody residues Y52_HC_ and D104_HC_. (G) I472, which is more than 8Å from the antibody surface, packs with the C480:C488 disulfide in the interior of the ACE2-binding ridge. Mutations to this residue may impact the conformation of this loop, which carries direct contact sites that escape antibody binding (shown as spheres), including residues Y473, A475, N487, and Y489.

**Table S1:** The mutation-level “escape-fraction” measured for each amino-acid mutation against each antibody. This CSV table is available at https://github.com/jbloomlab/SARS-CoV-2-RBD_MAP_clinical_Abs/blob/main/results/supp_data/REGN_and_LY-CoV016_raw_data.csv.

